# The effectiveness of prevention and early detection of tuberculosis in people living with HIV

**DOI:** 10.1101/2020.06.30.179895

**Authors:** Mikhail Sinitsyn, Evgeniy Belilovsky, Elena Bogorodskaya, Sergey Borisov, Dmitry Kudlay, Liudmila Slogotskaya

## Abstract

**Background:** The difficult epidemic situation caused by TB/HIV co-infection and, accordingly, the need to strengthen measures for the prevention of tuberculosis among HIV patients, identified the need to create a model of collaborating tuberculosis management facilities and AIDS centers.

**Study purpose:** analysis of the effectiveness of measures used in Moscow to prevent the spread of HIV/tuberculosis co-infection based on the developed new algorithm.

**Materials and methods:** The design of the study – a full-design retrospective cohort study of all HIV patients in tuberculosis preventive care and early detection unit for HIV patients (PCED TB Unit), which was organized in 2014 as a subdivision of Moscow TB Center on the premises of the Moscow AIDS Center. The study was conducted from 2014 to 2017. 22,190 HIV-infected patients were examined. In order to detect tuberculosis infection, all HIV patients were referred for skin testing with preparation Diaskintest, which represents recombinant proteins CFP10-ESAT6 and persons with a positive and doubtful reaction were examined for the diagnosis of tuberculosis and latent tuberculosis infection (LTBI). Preventive therapy (PT) was prescribed to patients with LTBI, which was also received by those with CD4^+^count ≤350 cells/mm^3^, regardless of the results of a skin test.

**Results:** Over a period of 4 years, Diaskintest was performed in 19,777 people. Positive results were in 857 patients (4.3%) [95% CI 4.1-4.6]. 131 patients were diagnosed with tuberculosis, the rest - with LTBI. The effectiveness of PT was estimated by the number of TB cases detected among HIV patients who completed PT (1757) and those who did not receive it (5990). The notification rate of TB in the first group was 228 [95%CI 65-582] per 100,000 patients, in the second 1486 [95%CI 1195–1825], which is 6,5 times higher. To assess the effectiveness of early detection of TB, the proportion of severity forms of pulmonary TB diagnosed in 87 patients at the PCED TB Unit using a new algorithm in 2016-2017 were compared with 411patients detected in different medical organizations of Moscow using conventional methods. The proportion of disseminated TB was lower in the first group (23.0% versus 44.5%) (p<0.01).

**Conclusion:** The study showed that the integration of the two services allows increases the accessibility of TB care and makes a significant contribution to improving the effectiveness of measures to prevent the TB spread among HIV patients. It supports the necessity of treating LTBI detected using the Diaskintest to prevent TB in HIV-infected patients. Significant effects that have been achieved include a reduction in the TB notification rate among HIV patients, improvement of the diagnostic structure of registered TB cases as a result of their timely detection.

## Introduction

HIV infection has a significant impact on the epidemiology of tuberculosis (TB) in the world and in the Russian Federation [1]. HIV-positive patients are 20–37 times more likely to have TB, as compared with HIV-negative individuals [2]. At the onset of latent tuberculosis infection (LTBI) in the human body, the likelihood of the disease is further increased, since the transition from latent TB to clinical manifestation is determined primarily by the immune status [3].

One of the strategies strongly recommended by the WHO (2018) for the eradication of tuberculosis worldwide is systematic testing that should allow detection and treatment of LTBI, in particular in HIV-infected people [4].

The World Health Organization (WHO) estimated in 1999 that 1.8 billion people, or one-third of the world’s population, were infected with *Mycobacterium tuberculosis* (Mtb) but without clinical symptoms of active tuberculosis (TB) which is the definition of latent TB infection (LTBI) [5]. In 2016 a WHO endorsed estimate updated the global prevalence of LTBI to 23% corresponding to 1.7 billion people infected worldwide [6,7]

Improved attention to LTBI screening and preventive treatment has been pointed out as crucial for The End TB Strategy for 2050 to be achieved [8,9]. It is hardly possible to eliminate TB unless progression to active TB is prevented underlining the need to determine the actual prevalence of LTBI and define hot spot areas [10].

The necessity of tuberculosis preventive treatment (PT) in HIV patients after exclusion of active tuberculosis is currently considered proven [11-14]

However, one of the main causes of the low effectiveness of measures to prevent the TB spread among HIV patients (spread of TB/HIV co-infection) is the lack of coordination between TB and HIV facilities [15-17]. The important question is which tests has the best potential for predicting the development of tuberculosis infection [14,18]

Many investigators believe that neither IGRA tests nor tuberculin tests have significant value in predicting the development of tuberculosis in individuals with positive results [14,19]. However, IGRA tests have a higher prognostic value in assessing the likelihood of the disease compared to tuberculin tests [19-22]. It has been shown that negative IGRA results are associated with a low probability of developing tuberculosis. Positive IGRA tests are more important than a tuberculin test in assessing the prognosis of tuberculosis in patients with LTBI combined with HIV infection [23,24]. Positive tests serve to identify individuals who should receive preventive anti-tuberculosis chemotherapy [25,26].

Recommended tests (tuberculin skin and IGRA tests) largely depend on the economic potential of the country, as a tuberculin test is significantly cheaper and easy to perform during screening, but has low specificity, particularly in BCG-vaccinated individuals [27-29].

IGRA tests, having high specificity, still have a number of serious disadvantages: high material costs, the need for an equipped laboratory, intravenous manipulation, which does not allow this method to be used for mass diagnostics [30-32].

The solution of the problem for Russia was the introduction of a skin test with CFP10 and ESAT6 proteins. In Russia, the medicinal product Diaskintest^®^ (manufactured by Generium according to the GMP standard) was developed, which is a complex of CFP10/ESAT6 recombinant proteins, produced by *Escherichia coli, BL21(DE3)/pCFP-ESAT*, intended for setting an skin test in a dose of 0.2 μg/0.1 ml according to the Mantoux test technique [33]. The test has been widely used in Russia since 2009 according to the order of the Ministry of Health of Russia.

Skin test with Diaskintest or recombinant tuberculosis allergen (RTA) demonstrated a high sensitivity (98%) in children with TB [34] 85% in adults and a high specificity (100%) [35-38]. Decreased sensitivity was observed in HIV patients with a CD cell count of less than 200 (22%) [39].

High effectiveness of preventive therapy has been demonstrated in patients with positive Diaskintest obtained during mass screening for tuberculosis in children [36,40] and in adults taking TNF-α inhibitors [41,42]

The introduction of Diaskintest affected the decrease in the incidence of tuberculosis in children and adolescents due to the better detection of patients with tuberculosis (TB), people with post-tuberculosis changes and LTBI and the use of preventive treatment for people at high risk groups. Detection of “minor” TB forms has improved [36,40]. A Danish skin test with C-Tb produced by Statens Serum Institut, which is similar to the Russian test, is now available. C-Tb sensitivity was similar in HIV-uninfected and HIV-infected patients 76.7% versus 69.5% and significantly diminished in HIV-infected patients with CD4^+^ cell count <100 [43-45].

The difficult epidemic situation caused by TB/HIV co-infection and, accordingly, the need to strengthen measures for the prevention of tuberculosis among HIV patients has determined the relevance of this work. It is necessary to determine the effectiveness of a model of collaborating tuberculosis management facilities and AIDS centers, including the mass screening for tuberculosis infection in HIV patients for several years and the use of preventive chemotherapy based on it.

### Study purpose

analysis of the effectiveness of measures used in Moscow to prevent the spread of HIV/tuberculosis co-infection based on the developed new algorithm of examination of people living with HIV.

### Objectives

1. Evaluation of the effectiveness of the early detection of TB based on the developed algorithm, by studying the severity of disease in new TB cases detected in HIV patients.
2. Assessment of the impact of PT on the TB incidence in patients with HIV infection.
3. Estimation of the LTBI spread among patients with HIV infection considered as an indicator of a potential risk of developing tuberculosis in this group of patients.
4. To evaluate the effectiveness of the new skin test using ESAT6-CFP10 proteins (Diaskintest) as an antigen for the mass screening of tuberculosis infection in patients with HIV and of the use of preventive therapy.

## Materials and methods

### Study designs

This study is designed as a full-design, open-label, retrospective cohort study of all HIV patients in TB preventive care and early detection unit for HIV patients (PCED TB Unit), which was organized in 2014 as a subdivision of the Scientific and Clinical Antituberculosis Center of Moscow Government Health Department (Moscow TB Center) on the premises of the Moscow City Center for the HIV/AIDS Prevention and Control (Moscow AIDS Center). The study was conducted from 2014 to 2017.

Moscow is a metropolis with a population of 12,7 million people.

Infectious disease physicians from the Moscow AIDS Center must refer all registered HIV patients to the PCED TB Unit for fluorography, consultation and skin testing with a recombinant tuberculosis allergen (Diaskintest).

#### Recombinant tuberculosis allergen (Diaskintest)

Recombinant tuberculosis allergen (Diaskintest) (CJSC GENERIUM, Russia) (manufactured according to the GMP standard) is a complex of CFP10/ESAT6 recombinant proteins, produced by Escherichia coli, BL21 (DE3)/pCFP-ESAT, intended for setting an skin test [33]. The test has been widely used in Russia since 2009 according to the order of the Ministry of Health of Russia.

Diaskintest was applied intradermally in a dose of 0.2 μg/0.1 ml according to the Mantoux test technique and transverse induration measured 72 hours later. When evaluating the response to the introduction, the test was regarded as positive in the presence of an induration of any size, doubtful in the presence of only hyperemia of any size, as negative - in the absence of induration or hyperemia

Persons with a positive or doubtful reaction to Diaskintest are subject to a thorough examination to identify active tuberculosis (pulmonary and extrapulmonary). They underwent a screening for clinical symptoms (cough, persistent fever, night sweats, weight loss), computed tomography (CT) of chest organs, ultrasound of other organs, and in the presence of changes in the chest organs – a sputum examination on Mtb culture and molecular genetic methods. In the presence of local changes in the chest, including the intrathoracic lymph nodes and the pleura, patients are subject to a assessment of changes in order to establish a diagnosis of tuberculosis. If a diagnosis of tuberculosis is detected, the patients are hospitalized in a 24–hour in-patient facility and given anti-tuberculosis therapy. Patients with positive reactions without local changes are subject to preventive therapy (PT). `Patients with a negative reaction to Diaskintest and` with CD4^+^count ≤350 cells/mm^3^, `are subject to the same examination as those with a positive reaction`. Further management strategy was determined in accordance with the “Algorithm for examination and management of HIV patients” (**Fig 1**).

**Fig 1.**
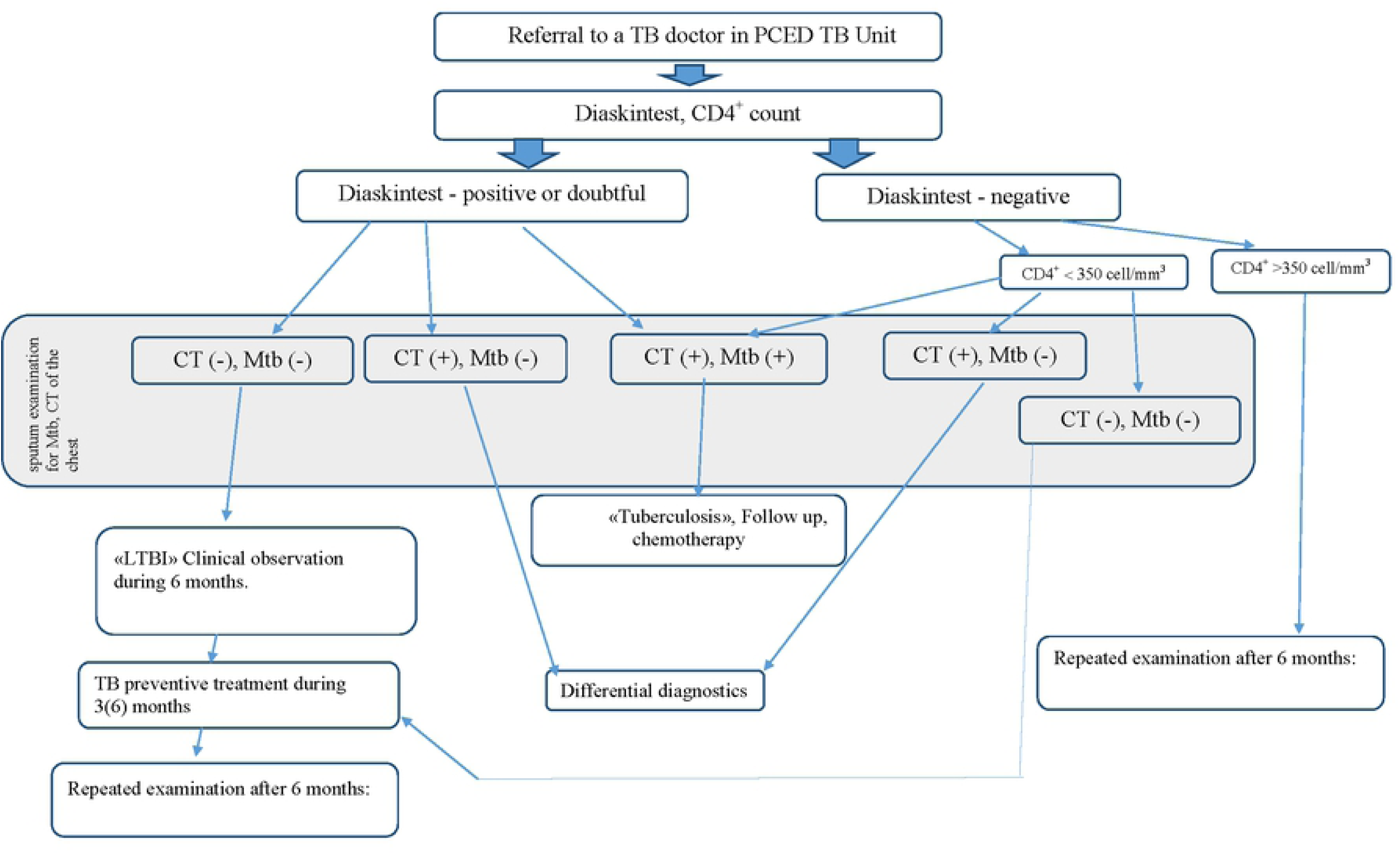
Algorithm for examination and management of HIV patients.

HIV-infected patients presenting with tuberculosis-related complaints and demonstrating abnormalities in the X-ray, patients with a history of contacts with tuberculosis-infected subjects or patients in whom differential diagnosis with other disorders was needed were referred by a specialist in infectious diseases for an unscheduled visit to a tuberculosis treatment specialist at the office PCED TB Unit. During the unscheduled visit to the tuberculosis treatment specialist, the following diagnostic tests were performed: a CT scan, Diaskintest, a sputum examination on Mtb culture and molecular genetic methods conducted in the central laboratory of Moscow TB Center, invasive tests with the purpose of morphological verification of the disorder (bronchoscopy, thoracoscopy, laparoscopy, lymph node biopsy). If these tests could not be performed in the outpatient setting, the tuberculosis treatment specialist referred the HIV-infected patient for hospitalization `in the clinic of infectious diseases (CID)`. A HIV-infected patient could be admitted to a tuberculosis clinic only after establishing the diagnosis of tuberculosis. In patients diagnosed with a latent tuberculosis infection, a tuberculosis treatment specialist prescribed PT and monitored the patient together with a specialist in infectious diseases for potential adverse reactions. The development of adverse events in patients receiving PT was monitored based on the evaluation of complaints and laboratory test results (mainly ALT and AST).

The following patients groups have been analyzed: 22 190 HIV-infected patients who were examined in PCED TB Unit) during 4 years (2014-2017), including 19 177 patients who were performed Diaskintest; 131 HIV-infected patients with new tuberculosis cases; 4413 HIV-infected patients who received preventive treatment (PT). To assess the effectiveness of PT, the cohorts of HIV patients, which visited the PCED TB Unit in 2015-2016 and underwent a full course of PT (1757), or not (5990) were analyzed. TB incidence was estimated based on new TB cases, which were registered during 1-2 years of follow up and no earlier than 12 months after the start of treatment, assuming that new TB cases detected earlier could be associated with tuberculosis not diagnosed before the start of PT.

To assess the effectiveness of early detection of TB, the forms of respiratory system tuberculosis revealed in 87 new TB cases among HIV-patients at the PCED TB Unit using a new algorithm in 2016-2017 were compared with 411 new TB cases detected in different medical organizations of Moscow using conventional methods.

#### Main approaches to the statistical analyses

Statistical processing was performed using the EpiInfo ™ software package [46]. All the statistical hypotheses used will be two-tailed, and p<0.05 will be used as the statistical significance level in all cases.

#### Ethical approvals

The trial was approved by the Ethics Committee of Scientific and Clinical Antituberculosis Center of Moscow Government Health Department, Moscow (number 3, 17.02.2018) conducted in accordance with the principles of Good Clinical Practice and the World Medical Association (WMA) Declaration of Helsinki adopted by the 18th WMA General Assembly, Helsinki, Finland, 1964 and subsequent amendments.

All parents give written informed consent for carrying out skin tests and any manner of examination and treatment. The patients gave their consent to the processing of the study data provided that no personal information is published. Access to medical documents for the conduct of this study was granted on 17.02. 2018.

## Results and discussion

### Results

A total of 22,190 HIV-infected patients were examined in PCED TB Unit. 131 (0.6%) [95% CI 0.5-0.7] HIV-infected patients were newly TB diagnosed with tuberculosis (6 per 1000 examined HIV-patients); 1282 (5.8%) [95% CI 5.5-6.1] cases required differential diagnosis, 84 patients (0.4%) [95% CI 0.3-0.5] had pulmonary lesions indicative of prior tuberculosis infection, which necessitated determination of its activity (Table 1).

**Table 1.** Characteristics of PCED TB Unit activities.

| Patient groups | Years |  |  |  |  |  |
| --- | --- | --- | --- | --- | --- | --- |
|  | 2014 (5 months) | 2015 | 2016 | 2017 | TOTAL | % |
| were examined (**total amount**), abs | 254 | 6968 | 7139 | 7829 | 22190 | 100 |
| including:<br>those who required additional tests, abs. | 40 | 233 | 462 | 547 | 1282 | 5.8% |
| new TB cases, abs. | 11 | 31 | 46 | 43 | 131 | 0.6% |
| demonstrated tuberculosis-related changes of unknown activity, abs. | 8 | 19 | 26 | 31 | 84 | 0.4% |

Over a period of 4 years, Diaskintest have been conducted in 20,942 HIV-infected patients. The tests were performed in 19,777 people (94.4%), were not conducted for medical reasons in 897 (4.3%) [95% CI 4.0-4.6] patients, and 268 patients (1.4%) [95% CI 1.2-1.5] refused to undergo the test. Positive results of Diaskintest were registered in 857 patients (4.3%) [95% CI 4.1-4.6], doubtful results – in 178 (0.9%) patients [95% CI 0.8-1.0], negative results – in 16,456 (83.2%) [95% CI 82.7-83.7]; 2,286 patients did not come for the test results assessment (11.6%) [95% CI 11.1-12.0] (Table 2).

**Table 2.** Results of Diaskintest in HIV-infected patients.

| Patient groups | Years |  |  |  |  |
| --- | --- | --- | --- | --- | --- |
|  | 2014 (5 months) | 2015 | 2016 | 2017 | Total |
| **Referred for testing, including:** | 229 | 6,011 | 7139 | 7,563 | 20,942 |
| Medical reasons for not conducting the test, abs. (%) | 20 (8.7) | 89 (1.5) | 232 (3.4) | 556 (7.4) | 897 (4.3) |
| patient's refusal, abs. (%) | 30 (13.1) | 64 (1.1) | 70 (0.9) | 104 (1.3) | 268 (1.4) |
| **Tests performed, abs. (%)** | 179 (78.2) | 5,858 (97.4) | 6,837 (95.7) | 6,903 (91.3) | 19,777 (94.4) |
| **including the result:** |  |  |  |  |  |
| positive, abs. (%) | 12 (6.5) | 257 (4.4) | 315 (4.6) | 273 (4.0) | 857 (4.3) |
| doubtful, abs. (%) | 1 (0.4) | 26 (0.4) | 58 (0.8) | 93 (1.3) | 178 (0.9) |
| negative, abs. (%) | 155 (86.6) | 4,873 (81.0) | 5,503 (80.5) | 5,925 (85.8) | 16,456 (83.2) |
| patients who did not come for the results assessment, abs. (%) | 12 (6.5) | 702 (11.9) | 961 (14.1) | 612 (8.9) | 2,286 (11.6) |

Among 131 new TB cases, 80 (61.1%) [95% CI: 52.2-69.5] had positive or doubtful reaction of the Diaskintest, and 51 patients (38.9%) [95% CI: 30.5-47.8] had negative reaction.

In 2016–2017, 89 HIV-infected patients were registered as a new TB cases. Respiratory TB was revealed in 87 patients, another TB localization - in two patients. Fluorography was performed in 65 patients with respiratory tuberculosis. In 30 (46.2%) cases [95% CI 33.7-59.0], it demonstrated changes that were typical for tuberculosis and in 5 (7.6%) [95% CI 2.5-17.0] cases - changes that were not typical for TB. It should be noted that in 30 patients (46.2%) [95% CI 33.7-59.0], fluorography did not reveal any abnormalities, while all 87 patients demonstrated changes in chest computed tomography (CT) scans.

To assess the effectiveness of early TB detection in HIV-infected patients, the number and proportion of different form in 87 new respiratory TB (RTB) cases, detected by PCED TB Unit using a new algorithm in 2016-2017 were compared with the forms 411 new RTB cases (HIV/TB-co-infected patients) in different medical facility of Moscow, using conventional methods in the same years (Table 3).

**Table 3.**
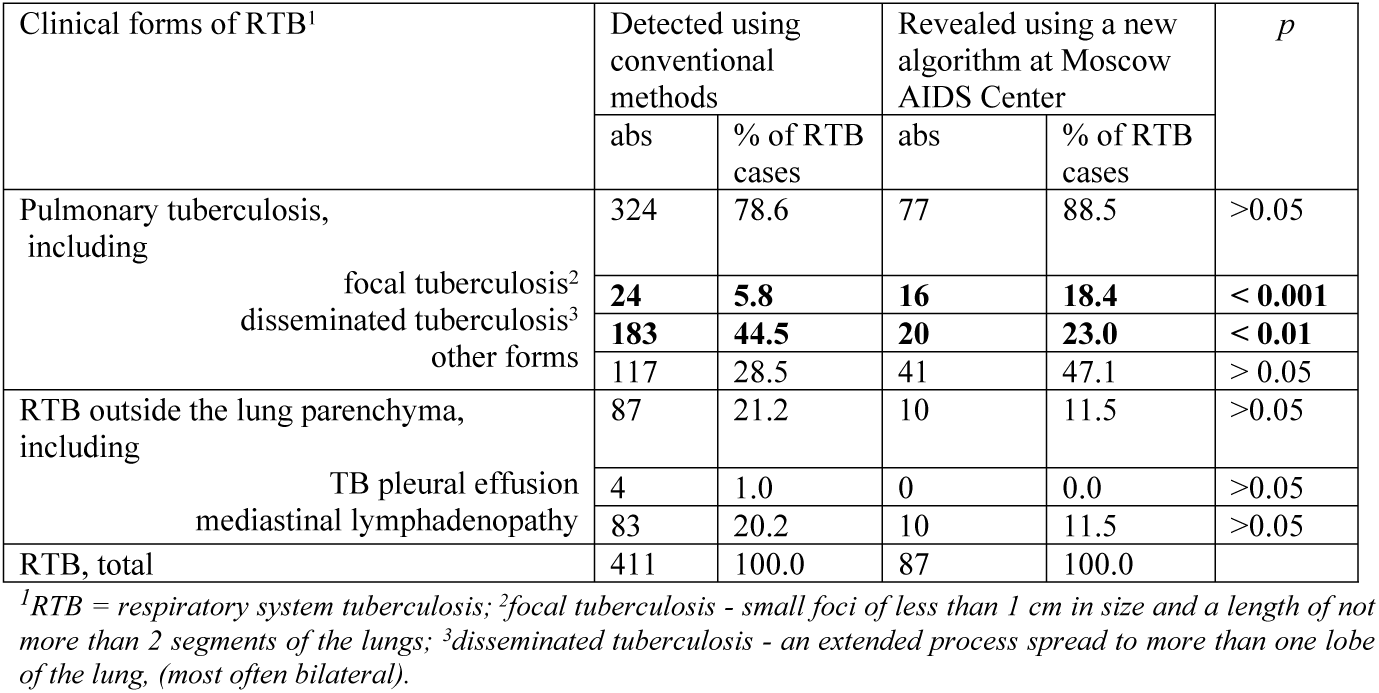
Clinical forms of respiratory system tuberculosis among new TB/HIV cases revealed by different approaches in 2016-2017.

| Clinical forms of RTB<sup>1</sup> | Detected using conventional methods |  | Revealed using a new algorithm at Moscow AIDS Center |  | *p* |
| --- | --- | --- | --- | --- | --- |
|  | abs | % of RTB cases | abs | % of RTB cases |  |
| Pulmonary tuberculosis, including | 324 | 78.6 | 77 | 88.5 | >0.05 |
| focal tuberculosis<sup>2</sup> | **24** | **5.8** | **16** | **18.4** | **< 0.001** |
| disseminated tuberculosis<sup>3</sup> | **183** | **44.5** | **20** | **23.0** | **< 0.01** |
| other forms | 117 | 28.5 | 41 | 47.1 | > 0.05 |
| RTB outside the lung parenchyma, including | 87 | 21.2 | 10 | 11.5 | >0.05 |
| TB pleural effusion | 4 | 1.0 | 0 | 0.0 | >0.05 |
| mediastinal lymphadenopathy | 83 | 20.2 | 10 | 11.5 | >0.05 |
| RTB, total | 411 | 100.0 | 87 | 100.0 |  |

<sup>1</sup>RTB = respiratory system tuberculosis; <sup>2</sup>focal tuberculosis - small foci of less than 1 cm in size and a length of not more than 2 segments of the lungs; <sup>3</sup>disseminated tuberculosis - an extended process spread to more than one lobe of the lung, (most often bilateral).

It has been demonstrated that the proportion of focal tuberculosis as a limited form in patients diagnosed as new RTB cases using the new algorithm in PCED TB Unit was significantly higher (18.4% versus 5.8%, respectively, *p*<0.001), while the proportion of disseminated tuberculosis (an extended process) was lower (23.0% versus 44.5%, *p*<0.01). The differences were even more evident for the pulmonary TB cases with higher proportion of focal tuberculosis (20.8% versus 7.4*%, p*<0,01), and lower proportion of disseminated tuberculosis (26% versus 56.3%, *p*<0.01).

Among all 89 new TB cases, “active TB detected” patients without tuberculosis-related complaints but detected by TB screening were registered in 75 (84.3%) [95%CI 75.0-91.1] cases; and 14 (15.7%) [95% CI 8.9-25.0] cases of tuberculosis were detected during unscheduled visits for evaluation of clinical symptoms. HIV/TB-co-infected patients were “actively” diagnosed (by screening) in other organizations in significantly lower proportion of cases - 50.9% (p<0.01).

#### Results and effectiveness of preventive treatment (PT) in HIV-infected patients

We analyzed the results of PT in 4,413 HIV-infected patients who had indications for it determined at the PCED TB Unit in 2014**-**2017 (Table 4).

**Table 4.** Tuberculosis preventive treatment in HIV-infected patients, n=4413.

| Years | Indicated | Contraindicated |  | Patient's refusal |  | Assigned |  | Completed |  | Interrupted |  |
| --- | --- | --- | --- | --- | --- | --- | --- | --- | --- | --- | --- |
|  | abs. (%) | abs. | % | abs. | % | abs. | % | abs. | % | abs. | % |
| 2014<br>(5 month) | 154<br>(100) | 6 | 3.9 | 31 | 20.1 | 117 | 76.0 | 60 | 39.0 | 57 | 37.0 |
| 2015 | 1060<br>(100) | 114 | 10.8 | 34 | 3.2 | 912 | 86.0 | 554 | 52.3 | 358 | 33.8 |
| 2016 | 1894<br>(100) | 152 | 8.0 | 19 | 1.0 | 1,723 | 91.0 | 1203 | 63.5 | 520 | 27.5 |
| 2017 | 1305<br>(100) | 98 | 7.5 | 13 | 1.0 | 1,194 | 91.5 | 978 | 74.9 | 216 | 16.6 |
| Total | 4413<br>(100) | 370 | 8.4 | 97 | 2.2 | 3,946 | 89.4 | 2795 | 63.3 | 1151 | 26.1 |

Over a period of 4 years, the proportion of patients receiving PT increased from 76.0% to 280 91.5%

In 2014 - 2017, 60 patients (39%) [95% CI 31.2-47.1], 554 (52.3%) [95% CI: 49.2-55.3], 1203 (63.5%) [95% CI: 61.3-65.7], and 978 (74.9%) [95% CI 72.5-77.3] respectively completed the course.

Antiretroviral treatment was used in 1581 (40.1%) of the total number of patients who received PT. The majority of HIV-infected patients receiving PT had low CD4^+^ cell counts. In 2014, 2015, 2016, CD4^+^ count <350 cells/µL was observed in 82 (70.1%) [95% CI 60.9-78.2], 768 (84.2%) [95% CI 81.7-86.5], 1627 (94.4%) [95% CI 93.2-95.5] and 1137 (95.2%) [95% CI 93.9-96.4] HIV-infected patients, respectively (Table 5).

**Table 5.** CD4^+^ cell counts in HIV-infected patients who received PT in 2014-2017, n=3946.

| CD4<sup>+</sup> count<br>(cells/mm<sup>3</sup>) | Years |  |  |  |
| --- | --- | --- | --- | --- |
|  | 2014 | 2015 | 2016 | 2017 |
| 0 – 99 | 18 (15.4%) | 107 (11.7%) | 369 (21.4%) | 290 (24.3%) |
| 100 – 349 | 64 (54.7%) | 661 (72.5%) | 1258 (73.0%) | 847 (70.9%) |
| ≥ 350 | 35 (29.9%) | 144 (15.8%) | 96 (5.6%) | 57 (4.8%) |
| Total | 117 (100%) | 912 (100%) | 1723 (100%) | 1194 (100%) |

Most commonly used PT regimens included two drugs (86%). Rifampicin-based treatment regimen was rarely used because of a rare potential adverse effect of its combination and antiretroviral drugs (n=153; 3.9%). Most patients received a combination of isoniazid plus pyrazinamide - 1,886 patients (34.3%), or isoniazid plus ethambutol - 1,353 patients (34.3%). Isoniazid monotherapy was used only in patients with co-morbidities who had contraindications for the use of rifampicin, pyrazinamide or ethambutol (Table 6).

**Table 6.** PT regimens for HIV-infected patients with LTBI, 2014-2017, n=3,946.

| Regimens | Years |  |  |  |  |  |
| --- | --- | --- | --- | --- | --- | --- |
|  | 2014 | 2015 | 2016 | 2017 | Total<br>abs., % | 95% CI |
| Isoniazid, 6 months, abs. | 77 | 155 | 190 | 132 | 554<br>(14.0%) | 12.9-15.2 |
| Isoniazid + ethambutol<br>3 months, abs. | 14 | 281 | 637 | 421 | 1,353<br>(34.3%) | 32.8-35.8 |
| Isoniazid + pyrazinamide,<br>3 months, abs. | 12 | 384 | 857 | 633 | 1886<br>(47.8%) | 46.2–49.4 |
| Isoniazid + rifampicin,<br>3 months, abs. | 14 | 92 | 39 | 8 | 153<br>(3.9%) | 3.3-4.5 |
| Total, abs. | 117 | 912 | 1,723 | 1,194 | 3,946<br>(100%) |  |

The safety and tolerability of different regimens used for prevention of tuberculosis in HIV-infected patients were analyzed. The analysis showed that 2795 of 3946 (70.8%) [95%CI 69.4–72. 2] patients in whom PT was prescribed successfully completed its course, while 1151 (29.2%) [95%CI 27.8–30.6] patients did not complete the course. Drug-induced adverse events were reported in 105 (2.7%) [95% CI 2.2-3.2] patients, and 1046 (26.5% [95% CI 25.1-27.9] patients made a decision not to continue treatment.

Monotherapy with isoniazid was inerrupted in 215 (38.8%) [95% CI 37.4-43.1] patients including 42 (7.6%) [95% CI 5.5-10.1] patients who were withdrawn from treatment for medical reasons (due to the development of adverse events) and 173 (31.2%) [95%CI 27.4-35.3] patients who made a decision not to continue treatment on their own (Table 7).

**Table 7.** PT regimens and PT outcomes in HIV-infected patients, 2014-2017, n=3,946.

| Regimens/<br>outcomes | 6H,<br>abs. (%) | 95%<br>CI | 3HE<br>abs. (%) | 95%<br>CI | 3HZ,<br>abs. (%) | 95%<br>CI | 3HR,<br>abs. (%) | 95%<br>CI | Total,<br>abs. (%) | 95%<br>CI |
| --- | --- | --- | --- | --- | --- | --- | --- | --- | --- | --- |
| **Completed treatment** | 339<br>(61.2) | 60.0-65.3 | 894<br>(66.1) | 63.5-68.6 | 1459<br>(77.4) | 75.4-79.2 | 103<br>(67.3) | 59.3-74.7 | 2795<br>(70.8) | 69.4-72.2 |
| **Did not complete treatment, including:** | 215<br>(38.8) | 34.7-43.0 | 459<br>(33.9) | 38.7-44.0 | 427<br>(22.6) | 20.8-24.6 | 50<br>(32.7) | 25.3-40.7 | 1151<br>(29.2) | 27.8-30.6 |
| due to adverse reactions | 42<br>(7.6) | 5.5-10.1 | 11<br>(0.8) | 0.4-1.4 | 37<br>(1.9) | 1.4-2.6 | 15<br>(9.8) | 5.5-15.6 | 105<br>(2.7) | 2.2-3.2 |
| interruption | 173<br>(31.2) | 27.4-35.3 | 448<br>(33.1) | 30.6-35.7 | 390<br>(20.7) | 18.9-22.6 | 35 (22.9) | 16.5-30.4 | 1046<br>(26.5) | 25.1-27.9 |
| **Total** | 554<br>(100) |  | 1353<br>(100) |  | 1886<br>(100) |  | 153<br>(100) |  | 3,946<br>(100) |  |

*Note: H – isoniazid, HE – isoniazid plus ethambutol, HZ – isoniazid plus pyrazinamide, HR – isoniazid plus rifampicin*

Among 1353 HIV-infected patients who received a combination of isoniazid plus ethambutol; 459 patients (33.9%) [95%CI 38.7-44.0] did not complete PT, including 448 patients (33.1%) [95%CI 30.6-35.7] who made a decision not to continue treatment. Adverse reactions leading to treatment interraption were reported in 11 (0.8%) [95% CI 0.4-1.5] among these patients.

Among 1,886 patients who received isoniazid plus pyrazinamide 427 patients (22.6%) [95% CI 20.8-24.6] did not comlete PT, including 390 (20.7%) patients [95%CI 18.9-22.6] who did it for unknown reasons and 37 (2.0%[95%CI 1.4-2.7]) who interrupted treatment due to adverse reactions. Among 153 patients who received a combination of isoniazid plus rifampicin; 50 (32.7%) [95% CI 25.3-40.7] patients stopped receiving PT, including due to drug intolerance in 15 cases (9.8%) [95%CI 5.6-15.6], while 35 patients made a decision not to continue this treatment (22.9%) [95%CI 16.5-30.4].

The frequency of adverse reactions in patients treated with isoniazid plus ethambutol for 3 months of daily was significantly lower than in patients receiving any of the other regimens (p<0.01). The frequency of adverse effects in patients receiving a three-month treatment with isoniazid plus pyrazinamide of daily was lower compared to isoniazid plus rifampicin combination (p<0.01).

The study demonstrated that a three-month PT of daily consisting of isoniazid plus ethambutol or isoniazid plus pyrazinamide was significantly better tolerated by HIV-infected patients compared to a 6-month treatment with isoniazid or a 3-month treatment with isoniazid plus rifampicin.

The frequency of treatment interraption was lower in patients treated of daily with isoniazid plus pyrazinamide or isoniazid plus rifampicin compared to those treated with isoniazid plus ethambutol or isoniazid monotherapy (p<0.01).

A total of 105 (2.7%) [95% CI: 2.2-3.2] patients developed adverse reactions leading to treatment interraption.

The effectiveness of PT in HIV-infected patients was assessed using the “Electronic Registry of HIV-infected patients followed up by the PCED TB Unit” We conducted a comparative analysis of the tuberculosis incidence rate in 7,747 patients who were followed up at PCED TB Unit in 2015-2016. This number included 1,757 patients who completed a course of PT, and 5,990 patients who did not receive PT during the same period.

To assess the effectiveness of PT, the TB notification rate was considered in the 3 following groups of HIV patients:

in group 1 - among 1757patients with HIV infection who completed a course of PT (1757 people),

in group 2 – among 5990 patients who visited PCED TB Unit in 2016 and to whom PT was not prescribed, or it was prescribed, but was interrupted (excluding those to whom PT was not assigned due to the fact that they had preventive therapy for the last two years), (5990 people),

in group 2a – among 1325 the same type of HIV patients as in the group 2, but whose CD4^+^count was ≤ 350 cells/mm^3^ (1325 people).

The effectiveness of PT was estimated by the number of new TB cases detected among HIV patients from group 1 after PT as of December 31, 2017. We considered only new TB cases registered one year after the date of appointment of PT. A total of 4 TB cases out of 1757 were recorded. Taking in account that time in risk of TB of these patients is 1-2 years, the TB incidence rate in HIV-infected patients who completed a course of PT was 227.7[95%CI 65-582] per 100 thousand patients with HIV infection.

For a more complete assessment of PT effectiveness, it should be taken into account that PT is performed in patients with an increased TB risk compared with other HIV patients, namely in patients with low CD4^+^ counts (mostly for CD4^+^≤350 cell/mm^3^.

Therefore, an estimate of the TB incidence in HIV patients was carried out both among patients who visited PCED TB Unit in 2016 and did not complete PT (group 2), and among the same patients with CD4+count ≤350 cells/mm^3^ (group 2a). New TB cases registered among these groups before 31.12.2017 were considered.

There were 89 new TB cases were recorded among 5990 examined patients from group 2 by December 31, 2017. Thus, the TB notification rate in this group of HIV patients was 1486 [95%CI 1195–1825] per 100 thousand of HIV patients.

Among 1325 HIV-patients with CD4+ ≤ 350 cells/mm^3^, who passed through PCED TB Unit, and who did not have or did not complete the course of PT, 33 new TB cases were registered by December 31, 2017, which determines the TB incidence estimation of 2491 [95%CI 1720–3480] per 100 thousand.

Thus, the analysis showed that the use of PT resulted in a 6.5-fold and 10.9-fold decrease for HIV-infected patients with low CD4+ count.

## Discussion

The effectiveness analysis of the algorithm that was developed for the detection and prevention of tuberculosis infection, which was organized on the basis of the combined work of the TB service and the AIDS center, revealed the following.

A total of 22,190 patients with HIV infection visited the PCED TB Unit over a period of 41 months (2014-2017). Among patients with HIV infection, 131 (0.6%) [95% CI 0.5-0.7] had tuberculosis diagnosed based on PCED TB Unit activities (6 per 1000 patients examined). Among these, 80 people (61.1% [95% CI 52.2-69.5]) had a positive or doubtful Diaskintest reactions.

We compared the effectiveness of various algorithms used for TB detecting in HIV patients by the PCED TB Unit in 2016-2017 and by other medical organizations in Moscow during the same period. In the former case, 89 patients were diagnosed with tuberculosis, including 87 with respiratory TB; in the latter instance, 411 patients were diagnosed. Comparison of the proportions of different clinical forms of tuberculosis in these groups showed that in the former case, the proportion of limited TB lesions (focal tuberculosis) was much higher (18.4% versus 5.8%, respectively) and that of disseminated tuberculosis was lower (23.0 % versus 44.5%), *p*<0.01.

Over a four-year period, Diaskintest was recommended in 20,942 patients with HIV infection (95.5%). The test was performed in 19,777 people (94.4%) and was positive in 857 people (4.3% [95% CI 4.1-4.6]), doubtful in 178 (0.9% [95% CI 0.8-1.0]), and negative in 16,456 (83.2% [95% CI 82.7-83.7]), while 2286 people did not come for evaluation (11.6% [95 % CI 11.1-12.0]).

It was shown that primary screening of tuberculosis using fluorography is less effective than primary screening using the skin test with Diaskintest. In particular, only 42.5% (95% CI: 31.0–54.6%) of patients, who subsequently had the TB diagnosis of tuberculosis confirmed, had changes characteristic of TB on fluorography. Esmail H et. al. showed that chest radiography (CXR) can be used to screen for evidence of subclinical TB, but is insensitive. They used ^18^F-FDG-PET/CT to identify pathology consistent with subclinical pulmonary TB in asymptomatic HIV-1 infected persons diagnosed with LTBI [13].

The Diaskintest showed a rather high sensitivity in 131 people who were diagnosed with tuberculosis, 62.4% (95% CI: 50.6–73.1%), which is consistent with the test results obtained in patients with tuberculosis and HIV infection using the Danish C-tb test, 69.5 % (95% CI 59.2– 78.5) [43-45].

All HIV-infected patients with positive or doubtful test results underwent a sputum test for *Mycobacterium tuberculosis* (bacteriological and molecular genetic), computed tomography, and, if necessary, invasive diagnostic tests to confirm the diagnosis, which allowed confirming or ruling out active tuberculosis in individuals with a positive Diaskintest and provide the identification of TB patients in the early stages of the disease [47].

After the differential diagnosis of tuberculosis and LTBI was carried out in subjects who had a positive Diaskintest reaction, all people with LTBI were prescribed preventive therapy.

Since HIV-infected patients with reduced CD4+ counts have an increased proportion of negative test results in the presence of tuberculosis infection, preventive therapy is prescribed when the number of these cells is less than 350 [48,49] even in patients with a negative Diaskintest reaction.

Preventive therapy administered to patients with HIV infection in accordance with the recommendations of the Moscow Society of Tuberculosis Physicians has shown its effectiveness and demonstrated the need to monitor for adverse reactions, as such monitoring allowed to obtain sufficiently high coverage and compliance. It was found that out of 3946 patients who were prescribed PT, 2795 people (70.8%[95% CI 69.4-72.2]) completed it successfully and 1151 (29.2%[95% CI 27.8-30.6]) did not complete the treatment course. Adverse events related to drug therapy were observed in 105 individuals (2.7%[95% CI 2.2-3.2]), while 1046 patients (26.5% [95% CI 25.1-27.9]) stopped taking the drugs on their own.

Two-drug PT regimens were used most commonly, in 86% of the cases. Rifampicin products were prescribed rather rarely due to a possible adverse combination with antiretroviral drugs, in 153 cases (3.9%). Most patients received regimens consisting of isoniazid plus pyrazinamide (1886 people, 34.3%) or isoniazid plus ethambutol (1353 patients, 34.3%). Isoniazid monotherapy for treatment was used only in patients who had contraindications to treatment with rifampicin, pyrazinamide, or ethambutol. A total of 105 patients (2.7% [95% CI 2.2-3.2]) developed adverse reactions that led to interraption of the treatment course. A three-month PT of daily consisting of isoniazid plus ethambutol or isoniazid plus pyrazinamide was significantly better tolerated by HIV-infected patients compared to a 6-month treatment with isoniazid or a 3-month treatment with isoniazid plus rifampicin. Sterling TR et al. have shown that 6 or 9 months of daily isoniazid are alternative recommended regimens; although efficacious, they have high toxicity risk and lower treatment completion rates, which decrease effectiveness [12]. As Esmail H. et al. have shown a proportion of HIV-1 infected adults with a negative screen for active TB (by sputum culture, CXR and symptom screen) and eligible for PT have evidence of subclinical disease hence it is likely that isoniazid monotherapy is often inadvertently prescribed [13].

The TB incidence rate estimation in HIV-infected patients was 6.5 times lower (p<0.01) in patients who received PT as compared with those who did not - 228 [95% CI 62-582] and 1486 [95% CI 1195-1825] per 100,000 patients with HIV infection respectively.

An organization in which primary screening of tuberculosis infection is carried out as close as possible to patients (on the premises of the AIDS center) by the tuberculosis service as a routine examination practice has proven to be effective.

Screening for tuberculosis infection using the Diaskintest is simple and cheap. It allows identification of the most vulnerable risk groups in the population. The prevalence of LTBI among HIV-infected people in Moscow was 4.5%, while in the general population it was 0.3-1.0% [50]. This examination method showed high efficiency in other risk groups, as preventive therapy in children and adolescents with positive results of the Diaskintest in a significant reduction in the incidence both in the risk groups and in the general population of the Moscow, where this method is used for the mass screening of children and adolescents every year [36,40].

Screening and monitoring of tuberculosis infection using the Diaskintest, followed by preventive therapy for patients with a positive test result, significantly reduced the number of patients with tuberculosis among those taking genetically engineered biological drugs, in particular, TNF-α inhibitors [41,42]. It was shown that in rheumatological patients Diaskintest can reduce indications for preventive therapy from 86% to 30-25%, as compared with the Mantoux test, at the initial screening stage and from 80% to 21% in the process of monitoring in subsequent months. After preventive therapy, cases of the disease were observed only in patients who did not receive it [41]. The use of prophylactic chemotherapy prevented the development of tuberculosis in all patients with inflammatory bowel disease who received treatment with TNF-α inhibitors, while 5% of patients who did not take preventive therapy developed tuberculosis. All patients developing the disease had a positive RTA test. The conversion to a positive RTA test in these patients increases the risk of tuberculosis 175 times [42].

It has already been acknowledged that IGRA tests and the new skin tests with the ESAT-6 and CFP-10 antigens conducted with the Russian Diaskintest and the Danish C-tb may offer additional benefits, primarily improved specificity, as compared with the PPD-based skin test [51,52]. It is emphasized that the availability of new tests, their cost-effectiveness and feasibility of use in the absence of resources, is a critical factor for expanding their scope of application [44].

## Conclusion

The study showed that the integration of the two services allows increases the accessibility of TB care and makes a significant contribution to improving the effectiveness of measures to prevent the TB spread among HIV patients. It supports the necessity of treating LTBI diagnosed using the Diaskintest to prevent TB in HIV-infected patients. Significant effects that have been achieved include a reduction in the TB incidence rate among HIV patients, improvement of the diagnostic structure of registered TB cases as a result of their timely detection.

## Acknowledgement

The study had no sponsorship.

## About authors

**Mikhail Sinitsyn**—Deputy Director, Clinical Research, Scientific and Clinical Antituberculosis Center of Moscow Government Health Department, Moscow; Russian Federation, Doctor of Medical Sciences.

Address: 10 Stromynka St., Moscow, 107014

Tel. +7 (499) 268-67-94

**Evgeniy Belilovskiy**—Head of the TB surveillance Department, Clinical Research, Scientific and Clinical Antituberculosis Center of Moscow Government Health Department, Moscow; Russian Federation, Ph.D., M.P.H.

Address: 10 Stromynka St., Moscow, 107014

Tel. + 7 (915) 190-90-10 ext. 203300

**Elena Bogorodskaya—**Director of the Clinical Research, Scientific and Clinical Antituberculosis Center of Moscow Government Health Department, Moscow; Russian Federation, Doctor of Medical Sciences.

Address: 10 Stromynka St., Moscow, 107014

Tel. + 7 (499) 268-00-05

**Sergey Borisov**— Deputy Director for Research and Clinical Work of the Clinical Research, Scientific and Clinical Antituberculosis Center of Moscow Government Health Department, Moscow; Russian Federation, Professor, Doctor of Medical Sciences.

Address: 10 Stromynka St., Moscow, 107014

Tel. + 7 (499) 785-25-14

**Dmitry Kudlay**—Leading Researcher, Laboratory of Personalized Medicine and Molecular Immunology, NRC Institute of Immunology FMBA of Russia, Doctor of Medical Sciences, Professor.

Address: 24 Kashirskoye shosse, Moscow, 115522.

Tel.: +7 (499)617-10-27

**Liudmila Slogotskaya** Head of the Clinical Research Department, Scientific and Clinical Antituberculosis Center of Moscow Government Health Department, Moscow; Russian Federation. Doctor of Medical Sciences

Address: 10 Stromynka St., Moscow, 107014

Tel. +7 (499) 268-04-15

## Author Contributions

**Conceptualization:** M. Sinitsyn, E. Bogorodskaya, D. Kudlay

**Formal analysis**: E. Belilovsky, L. Slogotskaya

**Investigation:** M. Sinitsyn, E. Belilovsky, E. Bogorodskaya, S. Borisov, D. Kudlay L. Slogotskaya

**Project administration**: M. Sinitsyn, E. Bogorodskaya, D. Kudlay.

**Writing ± original draft**: M. Sinitsyn

**Writing ± review & editing**: M. Sinitsyn, E. Belilovsky, E. Bogorodskaya, S. Borisov, D. Kudlay L. Slogotskaya

